# Risk of Seizures Induced by Intracranial Research Stimulation: Analysis of 770 Stimulation Sessions

**DOI:** 10.1101/688119

**Authors:** Hannah E. Goldstein, Elliot H. Smith, Robert E. Gross, Barbara C. Jobst, Bradley C. Lega, Michael R. Sperling, Gregory A. Worrell, Kareem A. Zaghloul, Paul A. Wanda, Michael J. Kahana, Daniel S. Rizzuto, Catherine A. Schevon, Guy M. McKhann, Sameer A. Sheth

**Affiliations:** Department of Neurological Surgery, Columbia University, New York, NY; Department of Neurology, Columbia University, New York, NY; Department of Neurosurgery, University of Utah, Salt Lake City, UT; Department of Neurosurgery, Emory University Hospital, Atlanta, GA; Department of Neurology, Dartmouth-Hitchcock Medical Center, Lebanon, NH; Department of Neurosurgery, University of Texas, Southwestern, Dallas, TX; Department of Neurology, Thomas Jefferson University Hospital, Philadelphia, PA; Department of Neurology, Mayo Clinic, Rochester, MN; Surgical Neurology Branch, NINDS, National Institutes of Health, Bethesda, MD; Department of Psychology, University of Pennsylvania, Philadelphia, PA; Department of Neurosurgery, Baylor College of Medicine, Houston, TX

**Author notes:** Corresponding author: Sameer A. Sheth, MD, PhD, Vice-Chair of Clinical Research, Department of Neurosurgery, Baylor College of Medicine, 7200 Cambridge, 9^th^ Floor, Houston, TX 77030.

**Keywords:** Research stimulation, seizures, intracranial stimulation, neuroethics

## Abstract

**Background:** Patients with medically refractory epilepsy often undergo intracranial electroencephalography (iEEG) monitoring to identify a seizure focus and determine their candidacy for surgical intervention. This clinically necessary monitoring period provides an increasingly utilized research opportunity to study human neurophysiology, however, ethical concerns demand a thorough appreciation of the associated risks.

**Objective:** We measured the incidence of research stimulation-associated seizures in a large multi-institutional study on brain stimulation’s effect on memory in order to determine if brain stimulation was statistically associated with seizure incidence, and identify potential risk factors for stimulation-associated seizures.

**Methods:** 188 subjects undergoing iEEG monitoring across 10 institutions participated in 770 research stimulation sessions over 3.5 years. Seizures within 30 minutes of a stimulation session were included in our retrospective analysis. We analyzed stimulation parameters, seizure incidence, and typical seizure patterns, to assess the likelihood that recorded seizures were stimulation-induced, rather than events that occurred by chance in epilepsy patients prone to seizing.

**Results:** In total, 14 seizures were included in our analysis. All events were single seizures, and no adverse events occurred. The mean amplitude of seizure-associated stimulation did not differ significantly from the mean amplitude delivered in sessions without seizures.

In order to determine the likelihood that seizures were stimulation induced, we used three sets of analyses: Visual iEEG analysis, statistical frequency, and power analyses. We determined that three of the 14 seizures were likely stimulation-induced, five were possibly stimulation-induced, and six were unlikely stimulation-induced. Overall, we estimate a rate of stimulation-induced seizures between 0.39% and 1.82% of sessions.

**Conclusions:** The rarity of stimulation-associated seizures, and that none added morbidity or affected the clinical course of any patient, are important findings for understanding the feasibility and safety of intracranial stimulation for research purposes.

## Introduction

Patients with medically refractory epilepsy, who comprise approximately one-third of epilepsy patients, often undergo an extensive workup attempting to identify the seizure onset zone and therefore determine their candidacy for surgical intervention. In cases where non-invasive elements of the workup do not adequately localize the seizure focus, invasive monitoring with intracranial electroencephalography (iEEG) may be clinically indicated. Examples of iEEG include subdural surface grids or strips, or intraparenchymal depth electrodes (stereo-EEG, or sEEG). After placement of iEEG electrodes in the operating room, seizure patients are monitored in an epilepsy-monitoring unit (EMU) as they are weaned off of their antiepileptic medications, in order to identify seizure foci. These patients, who are undergoing intracranial recordings for clinical purposes, provide an increasingly utilized opportunity to study human neurophysiology and understand the neurocircuitry underlying neurological and psychiatric disorders(1, 2). Intracranial stimulation in these patients for research purposes can provide unique and valuable information about both normal function and disease states. However, ethical concerns surrounding human-subjects research demand a thorough appreciation of the risks associated with such stimulation. One of the known risks of brain stimulation is the induction of seizures;(3-5) a risk that is increased in the epilepsy population, especially as patients are weaned off of their anti-epileptic medications in a controlled setting.

As human subjects research with brain stimulation becomes more common, a more thorough understanding of the safety profile and research-related risk of intracranial stimulation is paramount to our ability to safely and ethically conduct this research. It is important to determine safety parameters and potential thresholds for seizure induction in order to minimize associated risk. Furthermore, it is essential to determine the likelihood that seizures occurring during research sessions are stimulation-induced, or simply represent coincidental occurrences, when determining additional risk incurred by research participation.

In this study, we sought to characterize the nature and frequency of stimulation-associated seizures, looking at stimulation parameters, stimulation location, and baseline seizure patterns. We sought to assess potential stimulation safety thresholds, and better understand the overall risk associated with intracranial stimulation. We further sought to determine which seizures were more likely to be actually stimulation-induced, as opposed to coincidental occurrences in epilepsy patients prone to seizures. As stimulation-associated seizures are rare events overall, we aggregated data across a large multi-institutional cohort in order to achieve sufficient statistical power and generalizability.

## Materials and Methods

### Patient Selection and Stimulation Paradigm

The subjects for this study were patients with medically refractory epilepsy undergoing intracranial recording for seizure localization. All subjects had been enrolled in a multi-institutional research study (RAM, Restoring Active Memory), sponsored by the Defense Advanced Research Projects Agency (DARPA), whose goal was to enhance memory using intracranial stimulation. Subjects agreed to participate in research-related memory tasks with and without intracranial stimulation during their stay in the EMU. The continuous iEEG recording was monitored by an epileptologist during stimulation sessions in order to detect stimulation-associated after-discharge activity. The study was approved by each individual institution’s IRB and by the Department of Defense IRB.

The surgical procedure consisted of placement of either subdural grid/strip electrodes or sEEG electrodes, according to each center’s practice. High-resolution preoperative MRI and postoperative CT scans were obtained in all cases, using acquisition parameters standardized across all 10 centers. These imaging data allowed for accurate localization of electrode contacts within segmented MRIs to identify the exact position of each contact. Also part of the prospective design were instructions to each clinical site to document certain features of each stimulation-associated seizure: type of seizure (focal onset, aware/impaired aware, focal to bilateral tonic-clonic), whether its semiology was similar to the patient’s typical seizures, seizure severity, whether it required any additional measures such as respiratory support, and whether its occurrence increased overall length of stay in the EMU.

It was left to the discretion of each institution’s team to determine when during the EMU monitoring period to conduct research stimulation sessions – whether during the period of AED wean or only when on full AEDs, and whether before or after all seizures had been obtained for clinical purposes. Additionally, the exact location of research stimulation was determined by a combination of the central research team and each local clinical team. At the beginning of each research session, the team carried out a procedure to determine safe amplitudes for stimulation. This amplitude determination procedure consisted of two or three 1 s pulse trains ranging from 0.5mA up to the stimulation amplitude that would be used for the task. By protocol, stimulation was not to be carried out within 30 minutes of a preceding clinical seizure, and research sessions were terminated after the occurrence of a seizure, though not after the occurrence of afterdischarges, as long as the iEEG recording returned to baseline.

Clinical factors such as overall length of EMU stay, pattern of AED medication wean, etc. were also subject to each site’s discretion. The stimulation paradigm evolved over the course of the project, but overall parameters fell within the following range: anodic first balanced biphasic stimulation pulses of 300 μs per phase with amplitudes ranging between 0.1 and 3.5 mA, frequencies ranging between 10 and 200 Hz for an overall duration of 5 seconds, which yielded stimulation durations between 0.25 and 4.6 seconds.

### Data Analysis

At the outset of the DARPA RAM project, a “stimulation-associated seizure” was defined as any seizure occurring during or within 30 minutes of research stimulation. This was therefore an organic yet conservative threshold for inclusion in our retrospective analysis of seizure incidence. Seizure data recorded for any seizure occurring within 30 minutes of research stimulation included the type, duration, and severity of the seizure, the patient’s medication status, stimulation parameters used for that session, and baseline seizure information for the patient, when available. The prospective design of the study and the data coordination center ensured consistent data collection and annotation.

We first identified and collected data for all “stimulation-associated seizures.” The conservative operational definition of stimulation-associated seizures likely overestimated the actual rate of seizures caused by stimulation (“stimulation-induced seizures,”) because some of the seizures may have occurred spontaneously, as a consequence of the non-zero baseline seizure rate in this study population. In most clinical contexts, a seizure occurring more than five minutes, or sometimes even more than 30 seconds to 1 minute after delivered stimulation is not considered “stimulation-associated.” However, as the goal of this stimulation was not to trigger seizures, and as such, the occurrence of a seizure would be considered an adverse event, we decided to look at all seizures within the initially laid out 30 minute time window. We employed three further analyses to distinguish between coincidental seizures (spontaneous seizures around the time of stimulation) and those seizures likely to be induced by stimulation. The null hypothesis for these analyses was that all observed seizures were in fact stimulation-induced.

The first analysis was an electrographic analysis of the iEEG data to determine the temporal relationship between the time of stimulation and electrographic onset time of the seizure. A qualitative determination was then made as to whether the seizure could have been triggered by the delivered stimulation. Based on this analysis, seizures were determined to either be “concordant,” meaning the seizure clearly followed delivered stimulation in a temporal fashion, within a time window of 600 seconds, and therefore could have been triggered by stimulation; or “delayed,” meaning the seizure occurred greater than 600 seconds after stimulation delivery, but within the 30-minute time window parameter established at the start of the study. Given that any chosen time interval is necessarily arbitrary, 600 seconds was chosen as a reasonable and conservative one as it was felt that when assessing risk of research stimulation, it was better to err on the conservative end, rather than dismiss potential adverse events as unrelated.

We also determined the spatial relationship between stimulation delivery and seizure onset based on the contacts through which stimulation was delivered, and the contacts that showed earliest seizure onset. This information was not used to distinguish coincidental vs. stimulation-induced seizures, but rather to observe whether stimulation may induce, by our criteria, seizures at a distance from the stimulation site.

The second analysis was a statistical comparison of seizure frequency. We modeled each patient’s seizure rate with a Poisson distribution, which puts no upper limit on the total number of events, but assumes that they are relatively rare occurrences.(6) The parameters used to fit each patient’s individualized distribution were obtained from the number of seizures that occurred over his/her total EMU stay (poisson.test function in R). For each stimulation-associated seizure, we calculated the likelihood that it would have occurred by chance within the research session using the Poisson distribution. This analysis produced a p-value corresponding to the likelihood that the seizure that occurred during the stimulation session was not a chance occurrence. A significant p-value meant that the likelihood of seizure occurrence was outside the chance distribution, suggesting it was induced by stimulation. This hypothesis was tested for each of the 14 stimulation-associated seizures. The two-tailed threshold for statistical significance was set at p<0.05, with trend-level association defined as p<0.1. This analysis did assume that the frequency distribution of naturally occurring seizures remained constant throughout the EMU period. This assumption was imperfect, as AED doses and other factors were changing, but represents a reasonable initial estimate.

The final analysis for each seizure was designed to verify the statistical test for seizure frequency using a power analysis. We calculated a post-hoc power value for each of the aforementioned poisson tests. For example, if the previous analysis yielded a non-significant p-value (meaning the seizure fell within the expected background distribution) and had a high degree of statistical power (meaning there was enough data to have seen a significant difference had there been one), then the seizure was determined less likely to be stimulation-induced. Achieved power for each statistical test was calculated from effect size, type I error probability (alpha = 0.05), and the total sample size using R. Sufficient power was defined as > 0.8.

Using the above analyses, seizures were assigned to one of three categories based on the likelihood that the stimulation-associated seizure was in fact stimulation-induced as opposed to a coincidental occurrence. Categories were defined as follows: (1) likely stimulation-induced, (2) possibly stimulation-induced, and (3) unlikely stimulation-induced. The first category, likely stimulation-induced seizures, included any seizure with concordant iEEG data, and at least trend-level significance (p < 0.1) on rate analysis, regardless of the power analysis, or an insignificant p-value (p ≥ 0.1) on rate analysis, but with low power. The second category, possibly stimulation-induced, included seizures with concordant iEEG data but with insignificant p-values (p ≥ 0.1) with high power, or seizures for which the iEEG data was not available for full analysis. This was intentionally a conservative approach, meant to capture any event that could possibly be stimulation-induced, in order to determine an upper limit of actual stimulation-induced seizure rates.

The third category, unlikely stimulation-induced seizures, included seizures with delayed seizure onset on iEEG analysis, regardless of statistical significance. Again, the cut-off of 600 seconds between “concordant” and “delayed” on the iEEG analysis is somewhat arbitrary, as is the upper limit of 30 minutes established at the outset of the study. Arguably, both are conservative time frames, as in clinical electrical stimulation mapping, a seizure starting more than 10-30 seconds after stimulation delivery is generally thought to be incidental; however, in this study looking at the safety of stimulation for purely research purposes, these conservative estimations are purposefully meant to capture any seizure event that could possibly be related to stimulation **(Table 1)**.

**Table 1:**
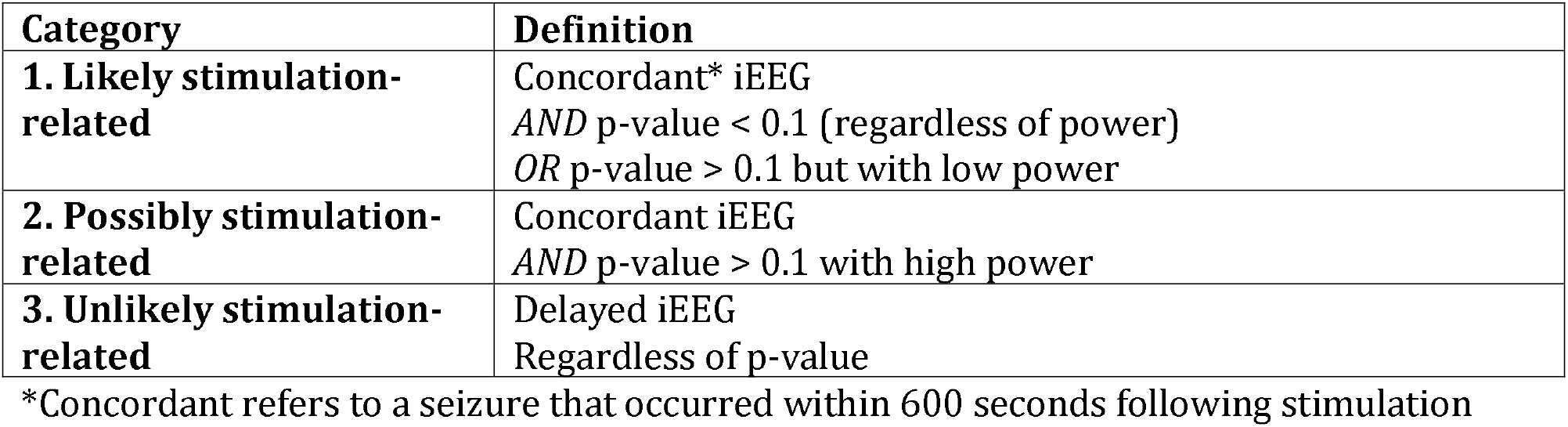
Stimulation-Relation Categories

| **Category** | **Definition** |
| --- | --- |
| **1. Likely stimulation-related** | Concordant* iEEG<br>*AND* p-value < 0.1 (regardless of power)<br>*OR* p-value > 0.1 but with low power |
| **2. Possibly stimulation-related** | Concordant iEEG<br>*AND* p-value > 0.1 with high power |
| **3. Unlikely stimulation-related** | Delayed iEEG<br>Regardless of p-value |

\*Concordant refers to a seizure that occurred within 600 seconds following stimulation

## Results

A total of 188 patients participated in 770 stimulation sessions over 3.5 years across 10 institutions. Defining a stimulation-associated seizure as any seizure occurring during research stimulation or within a subsequent 30-minute window, 14 seizures occurred in 12 different patients. The rate of seizures included in our study, which represents an upper limit of the true rate, was thus 1.82% of stimulation sessions.

Six of the 14 seizures (43%) were similar to the patient’s typical seizures in terms of semiology, onset, and spread. None of the seizures occurred in a cluster, with the majority (64%) being focal aware seizures. An additional 29% were focal impaired aware seizures; only one was a focal to bilateral tonic-clonic seizure. None of these stimulation-associated seizures required respiratory support beyond standard protocol-based placement of a nasal cannula. Per the assessment of the clinical team at each center, none of the stimulation-associated seizures were more severe than the patient’s typical seizure. The average clinical duration of the seizures was 79 seconds, with a range of 7-237 seconds. No adverse events occurred, and in the estimation of the clinical team at each site, these events did not affect overall length of stay in the EMU **(Table 2)**.

**Table 2:** Stimulation-Associated Seizure Characteristics

| **Seizure Characteristic** | **Number (%)**<br>**(total seizures = 14)** |
| --- | --- |
| Type of seizure |  |
| Focal onset aware | 9 (64%) |
| Focal onset impaired aware | 4 (29%) |
| Focal to bilateral tonic-clonic | 1 (7%) |
| Length of seizure | Mean: 79 seconds<br>Range: 7-237 seconds |
| More severe than patient's typical seizure? | 0 |
| Required intubation or respiratory support beyond NC? | 0 |
| Extended the amount of time the patient needed to be kept in the EMU? | 0 |
| Timing of seizure in relation to recording of all clinical seizures |  |
| Before | 9 (64%) |
| After | 5 (36%) |
| Patient on full home dose AEDs? |  |
| Yes | 2 (14%) |
| No | 11 (79%) |
| Unknown (not recorded) | 1 (7%) |
| Location of stim |  |
| CA1 | 4 (29%) |
| EC | 3 (21%) |
| Other | 7 (50%) |
| Location of stimulation relative to SOZ |  |
| Overlap in location of stim and SOZ | 2 (14%) |
| Overlap in location of stim and area of interictal spiking | 3 (21%) |
| No direct overlap | 9 (64%) |

In the majority of cases (64%) there was no overlap in the area of stimulation with either the known seizure onset zone or known areas of inter-ictal discharges. Of note, 11 of the 14 seizures (79%) occurred when patients were not on their full AED dose. In terms of stimulation parameters, there was no statistically significantly difference in stimulation amplitude between the sessions in which seizures occurred (mean 1.125 mA, range 0.25-2 mA) and the sessions without seizures (mean 1.06 mA, range 0.1-3.5 mA; p = 0.32). Current frequency and duration also did not differ between sessions without seizures and sessions with seizures.

In addition to the 14 seizures, there were 52 after-discharge (AD) events during the 770 stimulation sessions (6.8% of sessions), occurring in 36 different patients. Five of the patients who experienced AD events during stimulation sessions had seizures that were included in the current analysis. One of these seizures was preceded by three AD events that occurred in the same research stimulation session. Three other patients experienced ADs following stimulation with the same amplitude and frequency delivered at the same location in a previous stimulation session. Importantly, none of the stimulation-associated ADs evolved directly into seizures, meaning that on iEEG analysis, activity returned to normal after ADs, including in the sessions in which a seizure occurred.

We performed electrographic and statistical analyses in order to distinguish between coincidental seizures and those that were likely induced by stimulation. These 14 seizures are listed in **Tables 3 & 4** in the order that they occurred during the 3.5-year study. Looking first at the iEEG analysis, five seizures were found to be concordant, meaning that the seizure event occurred within 600 seconds following stimulation. Of these five “concordant” seizures, three were found to have trend-level significance on rate analysis, meaning there was a reasonable probability that they would not have occurred by chance, categorizing them as “likely” stimulation-induced. The remaining two of the concordant seizures had insignificant p-values on rate analysis, with high power, meaning based on the patients’ baseline seizure rates they were likely chance occurrences, pushing them into the “possibly” stimulation-induced category. Three additional seizures fell into the “possibly” stimulation-induced category, as their iEEG data were not available for review.

**Table 3:** Stimulation-Associated Seizures – Clinical Details

| Seizure | Location of Stim | Underlying Brain Pathology | Overlap with known SOZ or known areas of interictal discharges | Amplitude of Stim (mA) | Frequency of Stim (Hz) | Patient on full home dose AEDs | Type of Seizure | Duration of Seizure (seconds) |
| --- | --- | --- | --- | --- | --- | --- | --- | --- |
| 1 | R CA1 | None | No | 1.5 | 50 | No | Focal onset aware | 17 |
| 2 | R EC | None | Yes | 1 | 50 | No | Focal onset aware | 99 |
| 3 | R EC | None | Yes | 1.5 | 50 | No | Focal onset aware | 90 |
| 4 | L PHC | None | No | 0.5 | 50 | No | Focal onset impaired aware | 102 |
| 5 | L EC | None | No | 1.5 | 50 | Yes | Focal to bilateral tonic-clonic | 124 |
| 6 | L CA1 | None | No | 1 | 100-200 | No | Focal onset aware | 11 |
| 7 | R CA1 | None | Yes | 0.5 | 10-200 | No | Focal onset aware | 93 |
| 8 | R CA1 | None | Yes | 0.25 | 25 | No | Focal onset aware | 78 |
| 9 | R subiculum | None | No | 1 | 10-200 | No | Focal onset impaired aware | 146 |
| 10 | R superior frontal | R lateral frontal cortical dysplasia | Yes | 2 | 10-200 | No | Focal onset aware | 24 |
| 11 | R rostral middle frontal | R lateral frontal cortical dysplasia | No | 2 | 50 | No | Focal onset aware | 23 |
| 12 | R rostral anterior cingulate | None | No | 1 | 10-200 | Not recorded | Focal onset impaired aware | 7 |
| 13 | L MTG | None | No | 1 | 10-200 | No | Focal onset impaired aware | 237 |
| 14 | L6-7 | None | No | 1 | 200 | Yes | Focal onset aware | 57 |

(Stim = Stimulation; R = right; L = left; EC = entorhinal cortex; PHC = parahippocampal cortex; MTG = middle temporal gyrus)

**Table 4:** Stimulation-associated Seizures – Analysis

| **Seizure** | **iEEG Analysis** | **Rate analysis (p-value)** | **Power analysis** | **“Stimulation-associated” Category** |
| --- | --- | --- | --- | --- |
| 1 | Concordant | 0.052* | 0.96 | 1 – Likely |
| 2 | Concordant | 0.140 | 0.98 | 2 – Possibly |
| 3 | Concordant | 0.140 | 0.97 | 2 – Possibly |
| 4 | Delayed | 0.245 | 0.94 | 3 – Unlikely |
| 5 | Unavailable | 0.122 | 0.98 | 2 – Possibly |
| 6 | Unavailable | 0.120 | 1 | 2 – Possibly |
| 7 | Concordant | 0.097* | 1 | 1 – Likely |
| 8 | Delayed | 0.241 | 1 | 3 – Unlikely |
| 9 | Unavailable | 0.130 | 1 | 2 – Possibly |
| 10 | Delayed | 0.507 | 0.75 | 3 – Unlikely |
| 11 | Delayed | 0.507 | 0.75 | 3 – Unlikely |
| 12 | Delayed | 0.767 | 0.87 | 3 – Unlikely |
| 13 | Delayed | 0.090* | 1 | 3 – Unlikely |
| 14 | Concordant | 0.052* | 0.99 | 1 – Likely |

\*Denotes trend-level significance

The remaining six seizures occurred in a delayed fashion (after 600 seconds but within 30 minutes of stimulation) on iEEG analysis, so regardless of their rate analysis, they were categorized as “unlikely” stimulation-induced. Three of these seizures actually occurred after the completion of the stimulation session, though within the 30-minute time window, and so the seizures themselves were not captured on study-protocol iEEG, and were accordingly categorized as “delayed.” One seizure began electrographically before the stimulation was delivered during the session, but after the test stimulation for safety determination at the beginning of each trial. In this case, the seizure occurred more than 600 seconds after this test stimulation, placing it in the “delayed” iEEG analysis category, and ultimately in the “unlikely” stimulation-induced group.

The following section presents examples of each category of stimulation-associated seizure. A stimulation electrode and seizure onset electrode for each example are shown in Fig. 1A:

**Figure 1.**
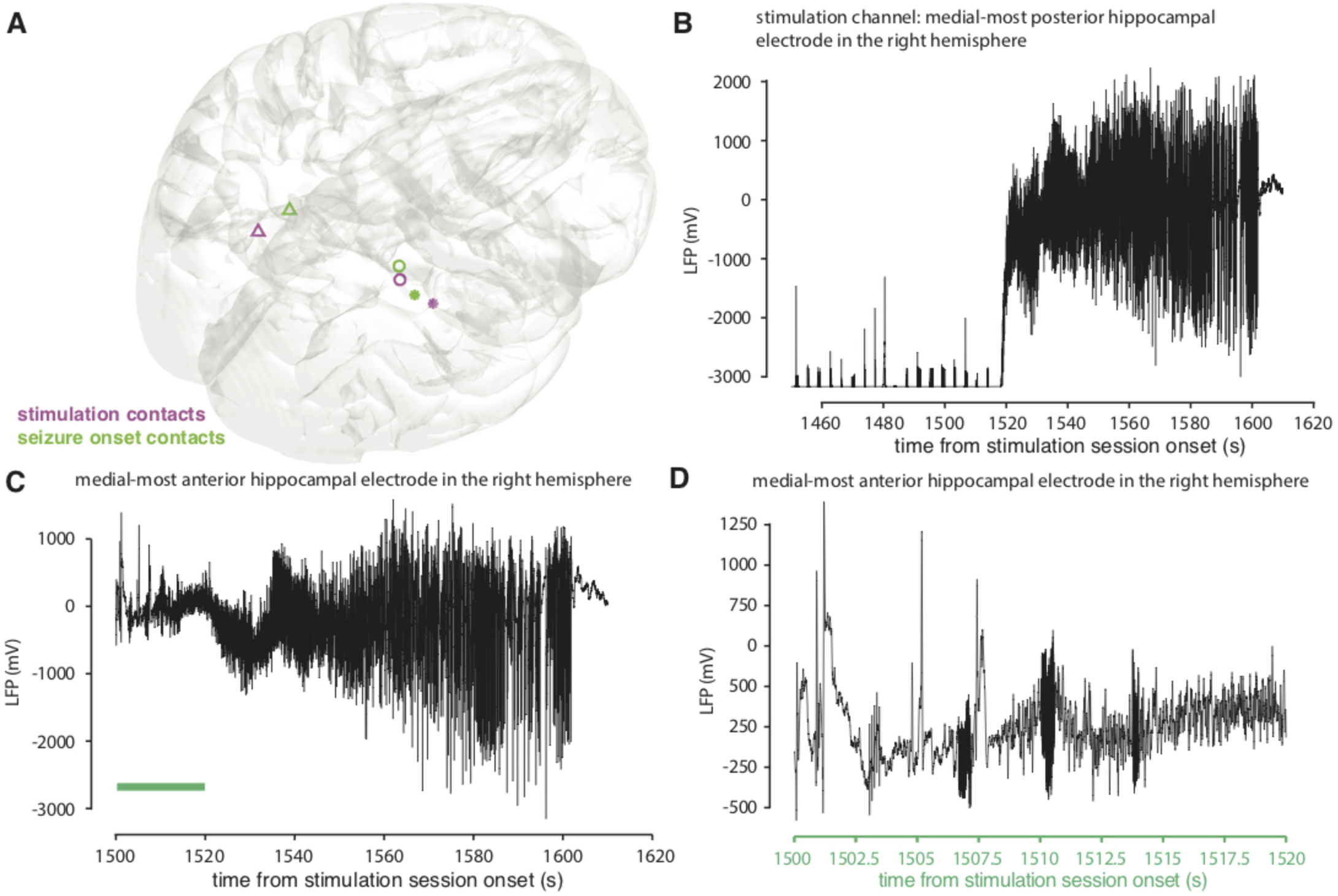
Seizure #7 (A) Localization of electrode channels in a 3D brain model; The open circles correspond to seizure #7, the asterisks to seizure #3, and the triangles to seizure #4. (B) iEEG recording from channel 66, the medial-most posterior hippocampal electrode contact, showing a train of stimulation current being delivered. This is followed by clear seizure activity. (C) Seizure activity can be seen most prominently in channel 57, which corresponds to the medial-most anterior hippocampal electrode contacts. (D) Expanded view of the recording from 1500 to 1520 seconds from panel C.

### Seizure #7: likely stimulation-induced

Seizure #7 was determined to be likely stimulation-induced. On iEEG analysis, a train of bipolar stimulation artifact can be seen in channels 65 and 66, which corresponds to the medial-most posterior hippocampal electrode contacts. Looking at channels 57 and 58, which correspond to the medial-most anterior hippocampal electrode contacts, there is clearly concordant seizure activity, as it begins while stimulation is still being delivered, followed by spread to the simulation channels and other surrounding electrodes **(Figure 1)**. The likelihood of causality based on seizure rate gave a p-value of 0.097, or trend level significance, with a power of 1. Based on iEEG analysis and rate analysis, this seizure was categorized as likely stimulation-induced.

### Seizure #3: possibly stimulation-induced

Seizure #3 is an example of a seizure that was determined to be possibly stimulation-induced. Looking at only the iEEG strip, stimulation delivered to the entorhinal cortex is followed by clear seizure activity, most prominent in the neighboring electrodes **(Figure 2)**. This seizure was clinically and electrographically similar to the other seizures the patient had while in the EMU. While based only on the iEEG analysis it is possible that the seizure was stimulation-induced, the rate analysis gave an insignificant p-value of 0.140, with a high power of 0.97, suggesting that this seizure could have been a chance occurrence. We therefore categorized this seizure as possibly stimulation-induced.

**Figure 2.**
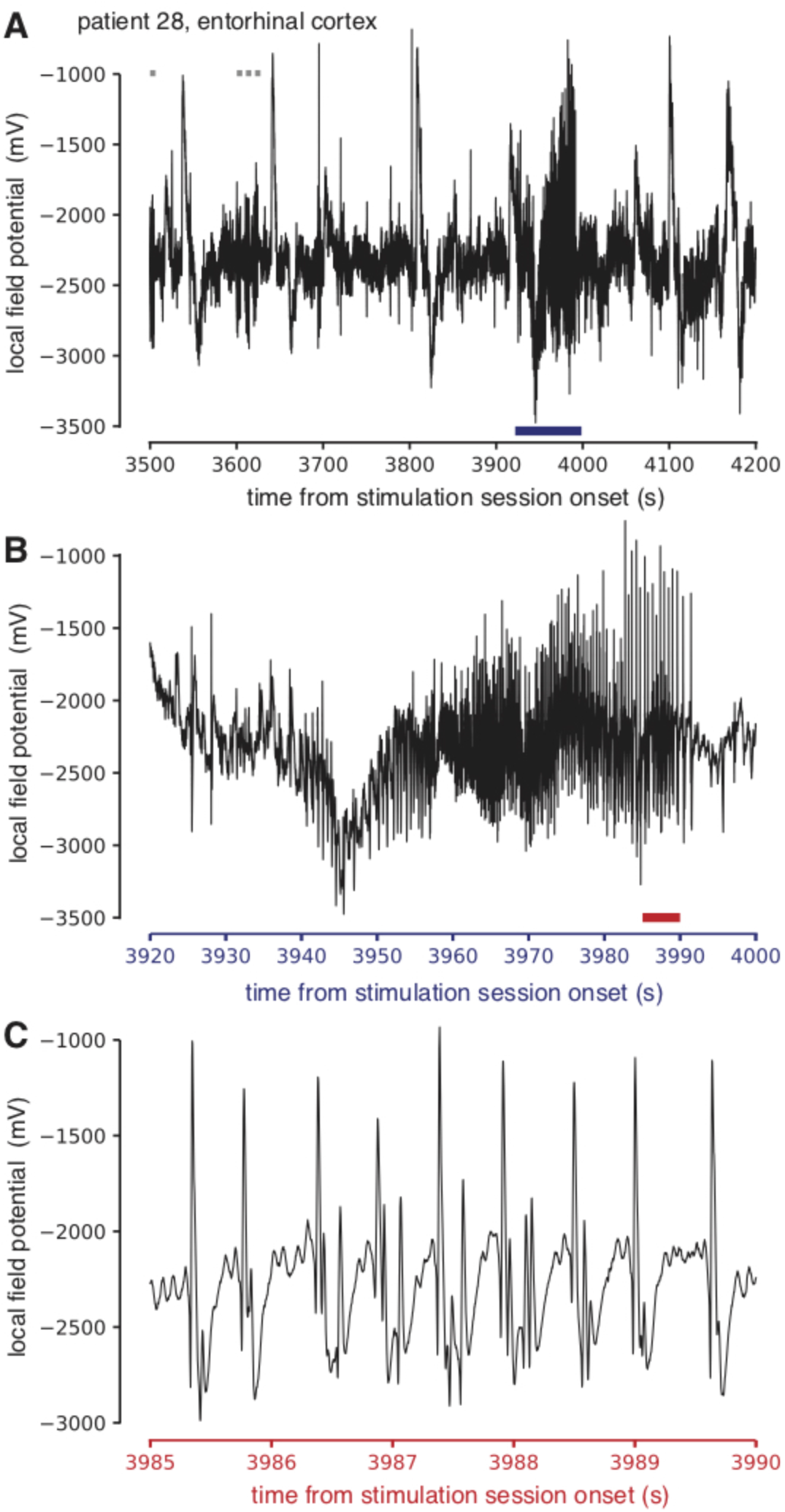
Seizure #3 (A) Stimulation delivered to entorhinal cortex, shown in gray bars along the top of the plot, followed by clear seizure activity in a neighboring channel (asterisks in Fig. 1A). (B) and (C) are expanded views of the seizure activity, which was both clinically and electrographically similar to the patient’s typical seizures.

### Seizure #4: unlikely stimulation-induced

Seizure #4 is an example of an unlikely stimulation-induced seizure. On iEEG, the seizure electrographically starts at approximately 520 seconds, just prior to stimulation delivery at 542 seconds in the posterior hippocampus **(Figure 3)**. In addition, the likelihood based on rate analysis was insignificant, with a p-value of 0.245 and a power of 0.94. The temporal order of seizure and stimulation on iEEG would make it seem impossible that stimulation could have produced this seizure. However, more than 600 seconds before this seizure, the researchers performed an amplitude determination test (see Methods for explanation of amplitude determination testing) consisting of two or three pulse trains at 0.5 mA. Because this short train of test stimulation preceded the seizure, we cannot definitively rule out a causal connection, even though the interval was >600 seconds. We therefore categorized this seizure as unlikely stimulation-induced to allow for this slim possibility.

**Figure 3.**
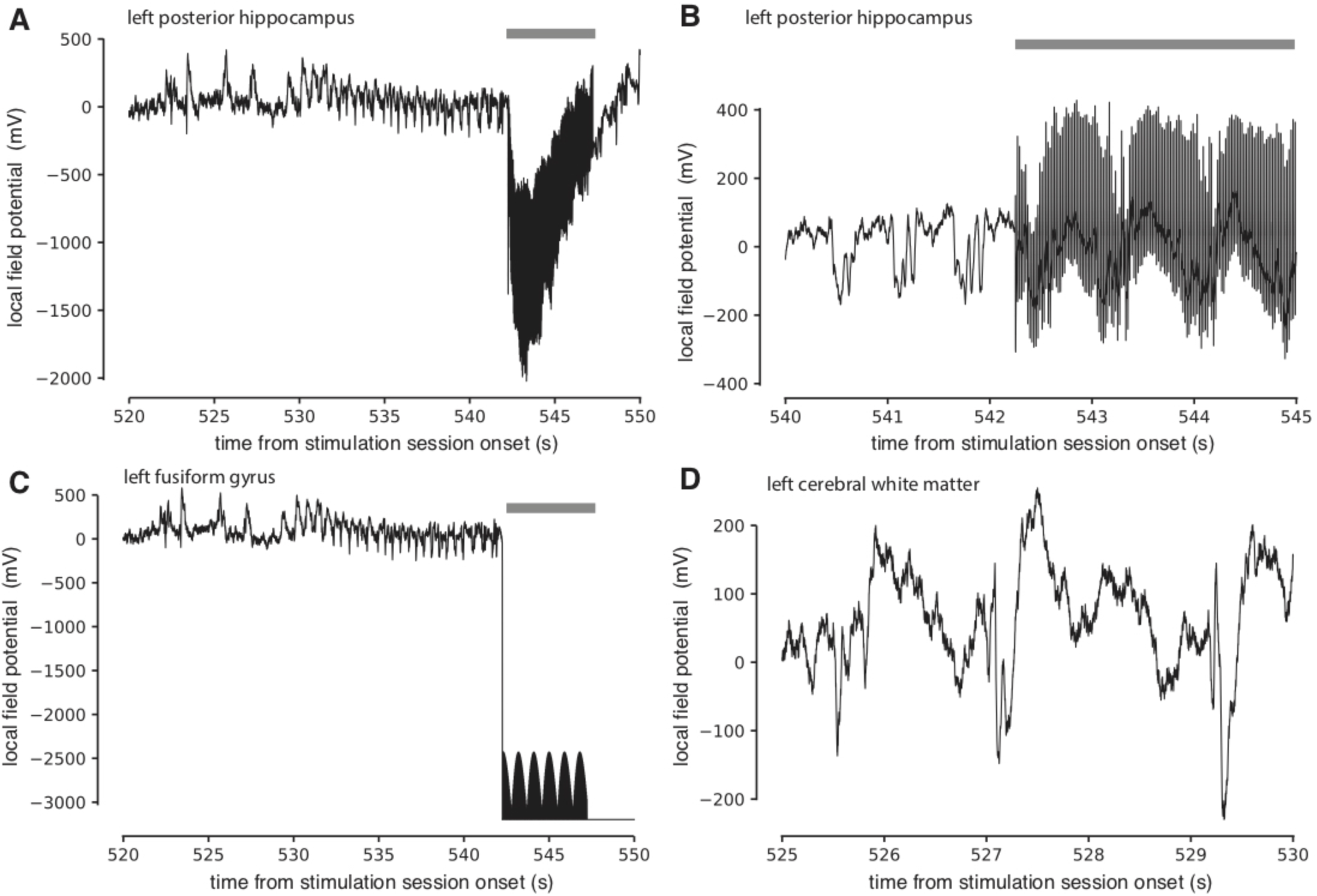
Seizure #4 (A, B) iEEG tracings from the stimulation channels in the posterior hippocampus (triangles in Fig. 1A) show seizure activity clearly starting prior to the onset of stimulation, which is delivered initially at approximately 542 seconds into the recording (indicated with gray bars along the top of the plot). (C) Overview, and (D) expanded view, captures of seizure activity, electrographically beginning just after 520 seconds, clearly before the onset of stimulation.

Putting the iEEG analysis, rate analysis, and power analysis together, three of the 14 seizures fell into the likely stimulation-induced category, resulting in an overall rate of 0.39% of all stimulation sessions. Five seizures fell into the possibly stimulation-induced category, and the remaining six seizures fell into the unlikely stimulation-induced category **(Table 4)**.

## Discussion

Based on our categorization method, three of the 14 seizures were determined to be likely stimulation-induced, five were determined to be possibly stimulation-induced, and the remaining six were determined to be unlikely stimulation-induced. These results suggest that “likely” stimulation-induced seizures occurred in only 0.39% of sessions. Inclusion of “possibly” stimulation-induced seizures results in a rate of 1.04% of sessions. A maximum rate of seizure occurrence of 1.82% of sessions results if the unlikely stimulation-induced seizures are included as well.

As the current study was conducted in patients with epilepsy, the majority of whom were not on their full dose AED medications during stimulation, one can infer that the seizure rate in this population is higher than in the general public. Therefore, as one extrapolates this data to future studies using intracranial stimulation or neuroprostheses in patients without epilepsy, one would expect to see an even lower incidence of stimulation-associated seizures. Adverse events associated with implanted neuroprostheses that have been reported include infection and device migration, without mention of seizures(7, 8).

Previous studies have reported seizures in 1.1 to 7.6% of patients during intra-operative stimulation, depending on the technique of stimulation utilized,(4, 9-21) and in up to 33-35% of patients during extra-operative cortical mapping.(3, 5) Importantly, both of these last studies reporting extra-operative stimulation-associated seizures employed higher stimulation amplitudes. Bank et al. report an initial current of 1-5 mA, which was increased by 1-2 mA per stimulation until the target maximum of 8-15 mA, while Aungaroon et al. report an initial current of 1-2mA, which was increased by 1-2 mA to a maximum current of 10 mA. Furthermore, while Aungaroon et al. report stimulation-associated seizures and stimulation-associated ADs separately, Bank et al. define a stimulation-evoked seizure “as after-discharges that evolved and were associated with clinical symptoms.” Neither study accounted for the possibility of coincidentally occurring seizures. Regardless, these numbers are still higher than the rate of seizures or after-discharges seen in stimulation sessions in the current study and may reflect the higher amplitudes used.

Kovac et al. investigated amplitude thresholds for inducing after-discharges, and report a mean stimulation intensity needed to elicit a clinical event of 3.3 ± 0.1 mA when using bipolar stimulation, and 3.9 ± 0.2mA when using monopolar stimulation.(22) Similarly, Karakis et al. report that an increase in stimulus intensity of 1 mA increases the likelihood of triggering intra-stimulation discharges, which are themselves associated with a 5-fold increase in odds of triggering after-discharges.(23) In our study, the mean amplitude used during stimulation sessions where a seizure occurred was 1.125 mA, and the maximum amplitude applied overall was 3.5 mA.

While we did not find a significant difference in maximum stimulation amplitudes between the sessions in which seizures occurred and the sessions without seizures, given the previous reports with higher rates of stimulation-associated seizures using higher stimulation amplitude, it appears that increased current amplitude is likely to trigger more seizures. Perhaps intuitively, the lowest possible stimulation parameters should be used in order to minimize risk of seizures during intracranial stimulation.

Four seizures occurred following stimulation parameters and locations known to trigger after-discharges. In order to reduce seizure risk further, one could avoid delivering stimulation if it has previously triggered an after-discharge at that location, regardless of whether the iEEG returned to baseline following the AD, or whether the AD occurred in a prior research session. Similarly, as 79% of patients were not on their full-dose home AED regimen when the seizure occurred, conducting research on epilepsy patients only when they are on full home-dose medications should intuitively reduce seizure frequency further. In our study, the location of stimulation did not appear to directly correlate with seizure occurrence. However, given that only 14 seizures occurred in our current study, it is possible that the sample size was too small to see a location effect.

While the size and multi-institutional nature of this study improves its generalizability, certain limitations to this study should be noted. For one, though criteria for what constituted a “stimulation-associated” seizure were clearly defined at the outset of the study, the reliance on individual reporting is subject to potential bias. One important element that was not closely controlled for was the effect of anti-epileptic medication wean on seizure frequency. We had access to rough measures of the stage of the medication wean at the time of each seizure, but not the exact doses relative to their pre-operative baseline, or medication status during stimulation sessions that were not associated with seizures. Overall, stimulation sessions were more likely to occur later during the patients’ EMU stays, and many centers waited until patients were back on their home dose of medication before carrying out stimulation sessions.

## Conclusion

In conclusion, while we know that seizures are a potential risk of research involving intracranial stimulation, the rate of stimulation-induced seizures identified in this large multi-institutional cohort was small. Importantly, none of the stimulation-associated seizures added morbidity or affected the clinical course of any of the patients. As intracranial stimulation is increasingly utilized to study normal and pathologic brain function, these results will be important for understanding the feasibility and safety of intracranial stimulation for research purposes.

## Abbreviations

AED: anti-epileptic drug
AD: after-discharge
EMU: epilepsy monitoring unit
iEEG: intracranial electroencephalography
sEEG: stereo-EEG

